# Identification of *cis*-regulatory modules responding to positional cues regulating the patterned expression of the *Drosophila* follicle cell fate determinants *mid* and *mirr*

**DOI:** 10.64898/2026.10.22.760784

**Authors:** S. De Vito, K. Ip, B. Rafanel, M. Fregoso Lomas, L.A. Nilson

**Author notes:** co-first author (equal contribution) (S. De Vito, K. Ip).

## Abstract

Using the patterning of the follicular epithelium of the Drosophila ovary as a model, we study how cells perceive spatial cues and how this information is interpreted by *cis*-regulatory regions of their downstream target genes.

During oogenesis, the oocyte secretes an inductive signal, Gurken (Grk), that activates the epidermal growth factor receptor (EGFR) within the overlying follicular epithelium. Grk/EGFR signaling outcome is context-dependent. At the posterior, Grk signals in the presence of the JAK/STAT pathway ligand Unpaired, inducing the posterior fate determinants *midline (mid)* and *H15.* At the anterior, in the presence of the BMP pathway ligand Dpp, Grk induces the dorsal anterior determinant *mirror (mirr)*. The choice between these targets is reinforced by mutual repression between Mid/H15 and Mirr, and by repression of *mirr* by Upd and *mid/H15* by Dpp.

To ask how these positional cues are interpreted, we screened non-coding regions from the *H15, mid* and *mirr* loci for CRMs that drive expression in patterns resembling that of the endogenous genes. We identify two regions each from the *mid* and *mirr* loci, each reproducing distinct aspects of their endogenous expression and regulation. These data provide a simplified system for asking how these sequences respond to upstream spatial cues.

**Summary statement:** Screening of non-coding sequences near Drosophila follicle cell fate genes identified *cis*-regulatory modules that respond to different subsets of known regulatory inputs and reproduce different aspects of their expression patterns.

## Introduction

During development, cells progressively acquire distinct gene expression profiles appropriate to their ultimate fate and function. These decisions often depend on extracellular spatial and temporal signals that regulate gene expression and confer upon a cell the properties appropriate for its position and developmental stage. The responses of individual cells to these spatial cues together generate dynamic patterns of gene expression within a developing tissue. How these upstream inputs are interpreted by regulatory elements of their target genes is not always clear.

Here we approach this question in the follicular epithelium of the Drosophila ovary, a well-characterized system for inductive signaling in epithelial patterning. The follicular epithelium surrounds individual germline clusters to form developmental units called egg chambers and, in late oogenesis, secretes the eggshell with clearly defined anterior-posterior (AP) and dorsal-ventral (DV) axes. Interestingly, these asymmetries are both established during oogenesis by the asymmetric localization of the EGFR ligand Gurken (Grk), which is secreted by the underlying oocyte [1, 2].

Early in development, the oocyte is localized at the posterior of the egg chamber, where secretion of Grk from the oocyte activates EGFR signaling in the overlying posterior follicle cells and induces expression of the T-box transcription factors (TFs) *midline* (*mid*) and its paralog *H15*, which determine posterior follicle cell fate [3, 4]. As the oocyte grows, Grk becomes localized to the dorsal anterior region where, instead of *mid/H15*, it induces expression of the homeodomain TF *mirror* (*mirr*), which determines dorsal anterior fate [5]. These observations show that the Grk/EGFR signal can induce expression of different target genes (*mid/H15* vs. *mirr*) depending on the context in which the signal is received.

We showed that the choice between these alternative Grk/EGFR signaling outcomes depends on additional localized extracellular signals. Posterior Grk/EGFR signaling occurs in the presence of Unpaired (Upd), a secreted JAK/STAT pathway ligand produced at the anterior and posterior poles of the epithelium [6, 7], and in this context induces *mid/H15* expression. Dorsal anterior Grk/EGFR signaling occurs in the presence of the BMP signaling ligand Decapentaplegic (Dpp), and in this context induces expression of *mirr* (Figure 1A, [8, 9]).

**Figure 1.**
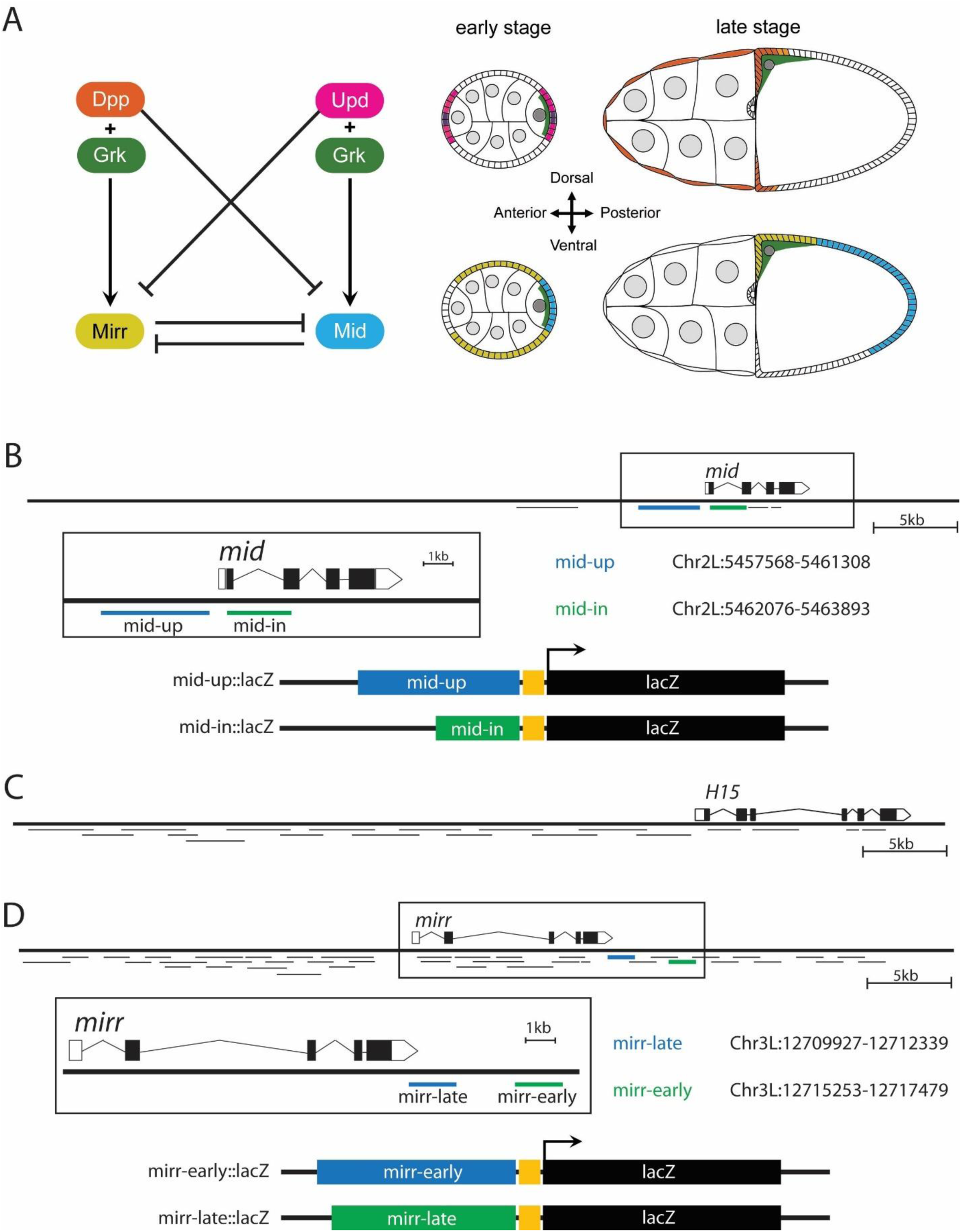
Enhancer screen for *mid* and *mirr*. (A) Model for regulatory network generating a bistable choice between induction of expression of the two alternative Grk/EGFR target genes *mid* and *mirr*. The EGFR ligand Grk (green) is secreted from the oocyte and localized at the oocyte nucleus (dark grey); Grk is localized at the posterior in early stages and at the dorsal anterior in later stages. The JAK/STAT signaling ligand Upd (magenta) is secreted by cells at the anterior and posterior poles of the tissue (purple), while the BMP pathway ligand Dpp is secreted by the anterior follicle cells, including the nurse-cell associated squamous “stretch cells” and the columnar “centripetal” cells (orange). In the presence of Upd, Grk/EGFR signaling pathway induces *mid* expression in posterior follicle cells (blue), while in the presence of Dpp Grk/EGFR signaling induces *mirr* expression in dorsal anterior follicle cells (yellow). (B-D) Gene schematics of the *mirr*, *mid*, and *H15* loci displaying genomic coverage of candidate regions included in the transcriptional reporters identified here (see also Table S1). Regions that drive follicle cell expression in a pattern reminiscent of the endogenous gene, and their coordinates (Assembly: BDGP Release 6 + ISO1 MT/dm6), are shown. Diagrams of the transcriptional reporters, in which these putative CRMs are placed upstream of an hsp70 basal promoter driving the lacZ gene, are also shown. (B) Two reporters, one containing genomic DNA upstream of the *mid* gene (R86G04, here referred to as mid-up) and the other containing genomic DNA from the first intron (R86F11, here referred to as mid-in) drive posterior expression. (C) No tested regions assigned to the H15 locusdrove reporter expression in posterior follicle cells. (D) Two reporters containing genomic DNA downstream of *mirr* produced patterns similar to the early (29814, referred to here as mirr-early) and late (29811, referred to here as mirr-late) components of the endogenous *mirr* expression pattern.

This choice of signaling outcome is reinforced by at least two known layers of negative regulation. One of these is reciprocal repression between *mid/H15* and *mirr*, reminiscent of double negative feedback loops that establish sharp gene expression boundaries in a range of developmental contexts [10]. Interestingly, independent of this mutual repression, *mid/H15* and *mirr* are each also repressed by the “opposing” positional cue: JAK/STAT signaling represses *mirr*, and Dpp signaling represses *mid* (Figure 1A). We have proposed that these elements define a bistable network that controls the choice between two Grk/EGFR targets [4], but how these multiple regulatory inputs are integrated, and the regions of the genome that ultimately interpret them, remains unclear.

To approach this question, we screened genomic sequences from the *H15*, *mid* and *mirr* loci to identify putative *cis*-regulatory modules (CRMs) that drive patterned expression of a transgenic reporter, then tested their response to the regulatory inputs that pattern endogenous *mirr* and *mid/H15* expression. For each locus, we identify two putative CRMs that drive reporter expression that exhibits key features of the expression pattern of the corresponding endogenous gene. Each putative CRM responds to a different subset of the identified regulatory inputs, suggesting that these signals are interpreted by genetically separable genomic regions. However, in the case of *mid*, reporters that concatenate the two identified regions exhibit an additive pattern of expression and regulation, suggesting that the larger genomic context is required for the integration of regulatory inputs that pattern the endogenous gene. This set of reporters therefore provides an interesting tool for investigating how upstream positional cues are interpreted by these CRMs to generate patterned gene expression, setting the stage for further mapping of these elements, for asking whether these elements are direct targets of the regulatory factors of this network, and for assessing the contribution of these elements to the regulation of their respective genes in the endogenous context.

## Results

### Identification of genomic sequences that drive patterned ovarian expression resembling that of *mid/H15* and *mirr*

To further understand how follicle cells integrate these multiple positional cues to generate mutually exclusive expression of these alternative Grk/EGFR target genes, we screened non-coding regions near the *mirr, mid* and *H15* loci for candidate CRMs that can drive reporter expression that recapitulates both their endogenous expression patterns and their responses to upstream positional cues.

We screened two genome-wide collections of transgenic transcriptional reporters [11–13], each consisting of 2-3kb of non-coding genomic DNA cloned upstream of a minimal promoter regulating the expression of the GAL4 transcriptional activator and inserted into the same genomic location [11–15]. We tested all available reporters assigned to the *mirr*, *mid*, and *H15* loci and selected those that drive patterned expression of a UAS-GFP transgene in the follicular epithelium (Table S1, Figure 1).

For each of these reporters, we characterized the reporters’ expression patterns and used mosaic analysis to test their responses to the upstream inputs that regulate the corresponding endogenous genes. Since some of these analyses utilize the GAL4-UAS system, e.g. as a clone marker or to drive knockdown or ectopic activation of signaling pathways, we generated lacZ versions of these reporters as well (see Materials & Methods). The expression patterns of these lacZ reporters, each inserted at two different target sites, closely resembles that of the original GAL4 version (Figure S1), suggesting that the transgenic construct and insertion site do not have a pronounced effect on the enhancer activities of these genomic sequences in the reporter context.

### Genomic sequences from the *mid* locus drive patterned expression resembling that of endogenous *mid*

For *mid*, two out of the five candidate reporters assigned to the *mid* locus produce expression at the posterior follicular epithelium, closely resembling that of endogenous Mid (Table S1, Figure 1B, Figure 2) [16].

**Figure 2.**
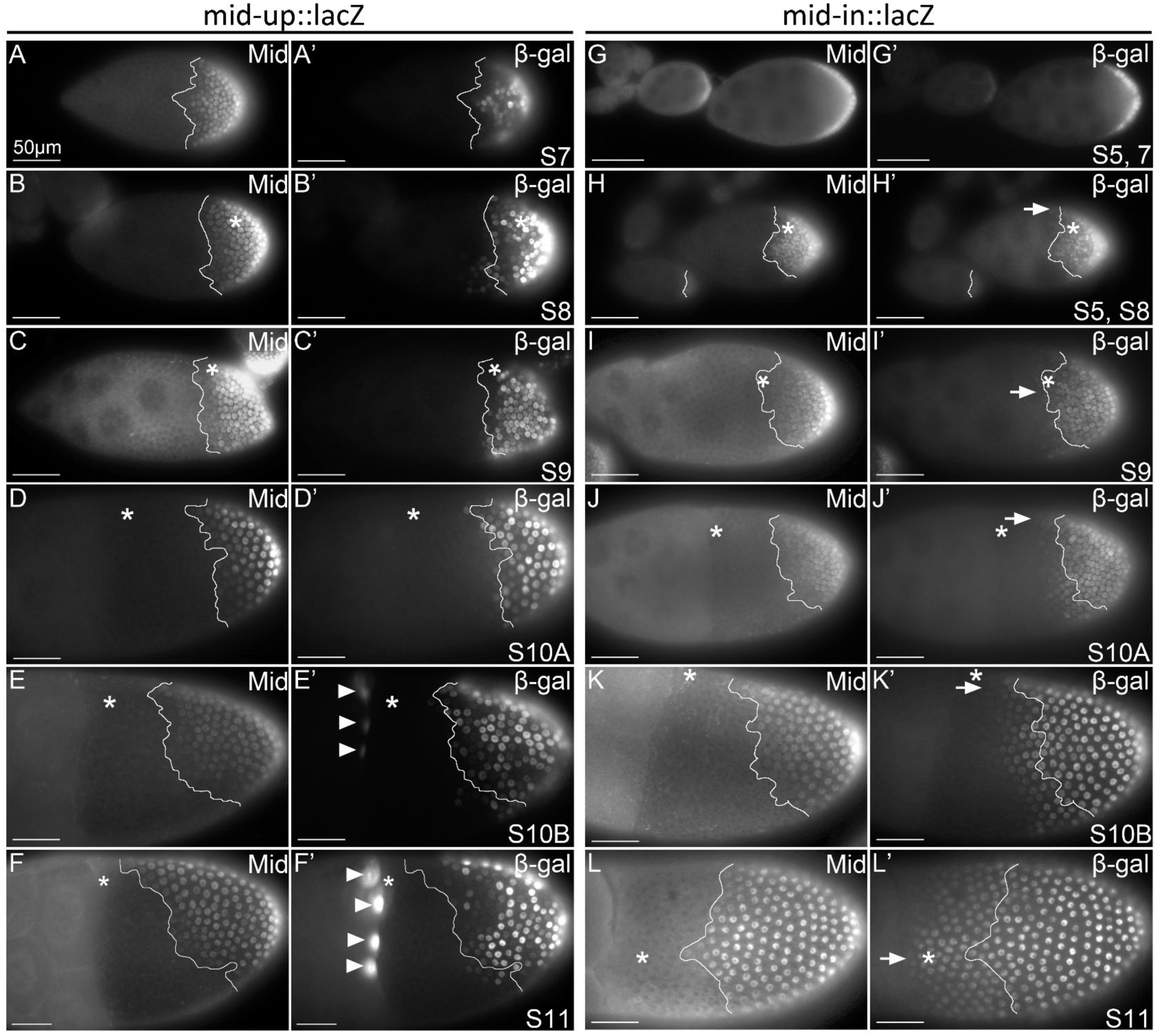
Expression of mid-up and mid-in LacZ reporters. Expression domains of mid-up and mid-in reporter in the follicular epithelium at different developmental stages (St7 – St11) with endogenous Mid expression pattern (white outlines) as comparison. Asterisks mark the dorsal side of the egg chamber. Scale bar = 50μm. (A-F’) mid-up drives reporter expression in the posterior columnar follicle cells in a similar pattern to endogenous Mid, albeit with some cell-to-cell variability in expression levels. At later stages, the dorsal peak of reporter expression is slightly less pronounced than that of endogenous Mid. Reporter expression is also observed in centripetal cells at the dorsal side (arrowheads). (G-L’) mid-in reporter displays an expression pattern that is posteriorly restricted in early stages but progressively expands anteriorly and ventrally such that it is expressed in all columnar follicle cells at St11. Arrows mark the anterior extent of reporter expression in columnar follicle cells. Mild expression is also seen in polar cell and centripetal expression (See Figure S1)

One of these (R86G04, which we refer to as mid-up) contains 3.6kb of genomic sequence ending 330bp upstream of the *mid* transcription start site. Expression of mid-up is first detectable at stage 7 of oogenesis, in a posterior pattern similar to that of endogenous Mid (Figure 2A-A’, Figure S1). At later stages, mid-up expression expands dorsally, generating a modest dorsal ventral bias that is reminiscent of, though less pronounced than, that of endogenous Mid [3, 4] (Figure 2B-F’). Within this posterior region, expression levels of mid-up are more varied among neighboring cells, compared to the more uniform levels of endogenous Mid. Unlike endogenous Mid, mid-up reporter expression is also detectable at stage 10B in centripetal cells at the dorsal side (Figure 2E-F’).

The second putative *mid* reporter (R86F11, which we refer to as mid-in) contains 1.8kb from the first intron of *mid*. At stage 7-8, like mid-up and endogenous Mid, mid-in reporter expression is confined to the posterior domain (Figure 2G-H’,[16]). However, this expression progressively expands anteriorly over the course of oogenesis beyond the endogenous Mid expression domain, and by stage 11 reporter expression is seen in all columnar follicle cells (Figure 2I-L’). mid-in also drives weak reporter expression within the border cell cluster and in some centripetal cells, where expression of endogenous Mid is not detectable (Figure S2).

*H15* and *mid* are paralogs separated by only 45 kilobases, and exhibit overlapping patterns of posterior expression [3]. For H15, none of the 19 candidates assigned to the *H15* locus produced patterned expression in the follicular epithelium (Figure 1C, Table S1). While the elements that regulate posterior expression of *H15* could lie elsewhere in the genome, the proximity of *H15* and *mid* loci and their similar expression patterns suggest that they might share *cis*-regulatory elements. Indeed, a recent study showed that R86G04 contains *cis*-regulatory elements that regulate *H15* expression [16]. We therefore focused on the genomic regions attributed to the *mid* locus.

### Genomic sequences from the *mirr* locus drive patterned expression resembling that of endogenous *mirr*

For *mirr*, we tested each of the 40 candidate reporters assigned to the *mirr* locus and found two that drive patterned expression in the follicular epithelium (Figure 1D, Figure 3, Table S1). Neither recapitulates the entire spatiotemporal pattern of the endogenous *mirr* expression, as revealed by *mirr^l(3)cD1^* enhancer trap, whose expression matches that of endogenous *mirr* (Figure S3)[5]. Instead, each appears to capture independent elements of the expression pattern.

**Figure 3.**
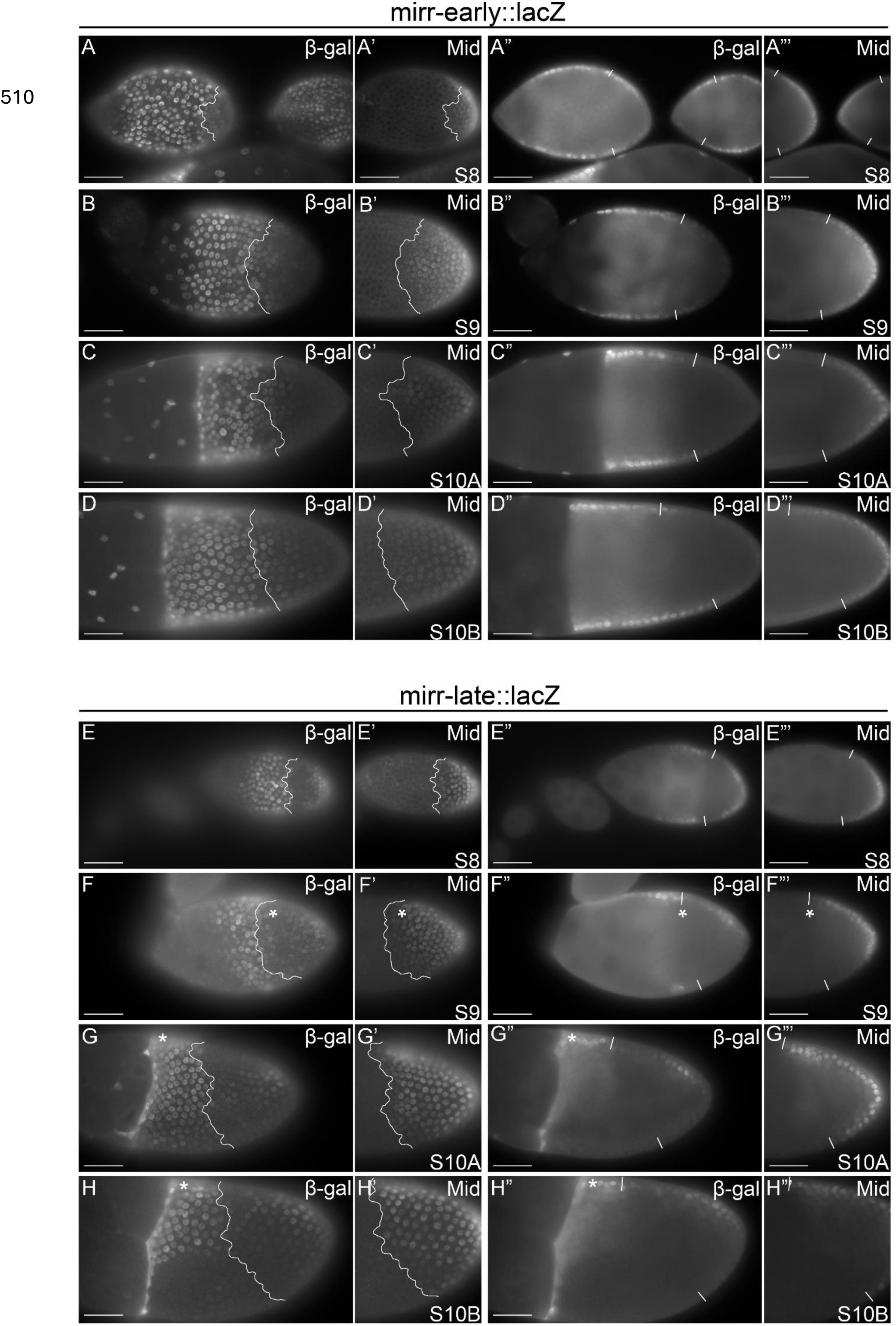
Expression of mirr-early and mirr-late LacZ reporters. Expression domains of mirr-early and mirr-late reporters in the follicular epithelium at different developmental stages (St8 – St10B) with endogenous Mid expression pattern as comparison (white outlines). Asterisk marks the dorsal side of the egg chamber where applicable. Left columns shows surface view; right columns show the cross-sectional views of the same egg chamber. Scale bars = 50μm. (A-D”’) Throughout development, mirr-early reporter expression is excluded from the anterior terminal domain but present in the main body follicle cells and present at a low level in posterior terminal domains. Mirr-early reporter expression is present in the Mid expression domain and does not display dorsal/ventral bias at later developmental stages. (E-H”’) mirr-late reporter drives expression at the main body follicle cells and at a group of posterior follicle cells. The posterior expression boundary of the main body domain is complementary to that of Mid expression domain (white line); the posterior domain is dorsally biased. mirr-late reporter is also expressed in the posterior follicle cells. Beginning stage 9, reporter expression in both the main body and posterior domains displays a dorsal (asterisk bias ). Beginning stage 10B, reporter expression is present at centripetal cells.

One reporter (VT29814, which we refer to as mirr-early), contains 2.2kb of genomic DNA beginning ∼5kb downstream of the *mirr* transcription termination site. The mirr-early reporter is expressed at early stages (8-9) in most follicle cells but is absent or reduced at the anterior and posterior terminal domains (Figure 3A-B”’). The resulting pattern of “mainbody” expression resembles that of endogenous *mirr* at these stages (Figure S3). [5] Interestingly, unlike the *mirr^l(3)cD1^* enhancer trap, the posterior limit of mirr-early reporter expression overlaps slightly with the anterior limit of the Mid expression domain and does not display dorsal/ventral bias at later developmental stages (Figure 3C-D”’).

The second reporter (VT29811, which we refer to as mirr-late) is enriched in later stages in the dorsal anterior region and in the ventral belt domain of the epithelium (Figure 3E-H”’). This pattern resembles that of endogenous *mirr* at these stages, though without the dorsal anterior high point observed for endogenous *mirr* (Figure S3)[5]. Unlike endogenous *mirr*, however, the mirr-late reporter also exhibits expression from stage 8 in the posterior domain, with a clear dorsal bias by stage 10A.

The observation that these genomic regions drive reporter expression resembling different components of the endogenous *mirr* pattern suggests that the dynamic pattern of endogenous *mirr* expression is controlled by different *cis*-regulatory elements. The novel posterior expression of mirr-late reporter suggests that its sequences contain enhancer activity that is masked in the endogenous genomic context.

### Assessing the response of *mid* and *mirr* reporters to the positional cues that drive patterned expression of endogenous *mid* and *mirr*

The observation that these non-coding regions from the *mirr* and *mid* loci drive patterned reporter expression resembling features of the endogenous *mirr* and *mid* genes suggests that they contain *cis*-regulatory elements that respond to the positional cues that regulate *mirr* and *mid* expression in this tissue. As discussed above (see Figure 1A), these include extrinsic spatial information from the localized ligands Upd and Dpp, which activate the JAK/STAT and Dpp signaling pathways, respectively [4, 6, 7, 9, 17, 18], as well as intrinsic information in the form of the *mirr*/*mid* mutual repression [4]. To test this hypothesis, we used the Flp-FRT system [19, 20] to clonally manipulate the activity of these regulators, using loss-of-function alleles or ectopic expression using the GAL4/UAS system [21]. We induced ectopic activation of the JAK/STAT and Dpp signaling pathways through clonal overexpression of the <u>JAKJanus</u> kinase Hopscotch (Hop) [22] or clonal expression of a constitutively active form of the type I BMP receptor Thickveins (*tkv*) [23], respectively. An effect on reporter expression <u>that</u> mimics that observed for the endogenous gene would suggest that the genomic sequence contained in the reporter includes one or more elements that respond, directly or indirectly, to these upstream inputs. The expression of endogenous Mid serves as internal control for the clonal analysis (Figure S4, Figure S5).

As detailed below, for both *mid* and *mirr,* the responses to these multiple regulatory inputs are distributed across the reporters we identified, roughly mapping the response elements to separable genomic regions.

### mid-up responds to Upd and Mirr, but not Dpp

Since endogenous Mid expression is positively regulated by Upd, and mid-up reporter is expressed in a pattern that closely matches that of endogenous Mid, we hypothesized that mid-up is also positively regulated by Upd. Indeed, we found that mid-up reporter expression is lost in cells homozygous for an amorphic allele of *StatS2E* [17] (Figure 4B-B’), with a penetrance comparable to that of endogenous Mid (Figure S4A’)[4].

**Figure 4.**
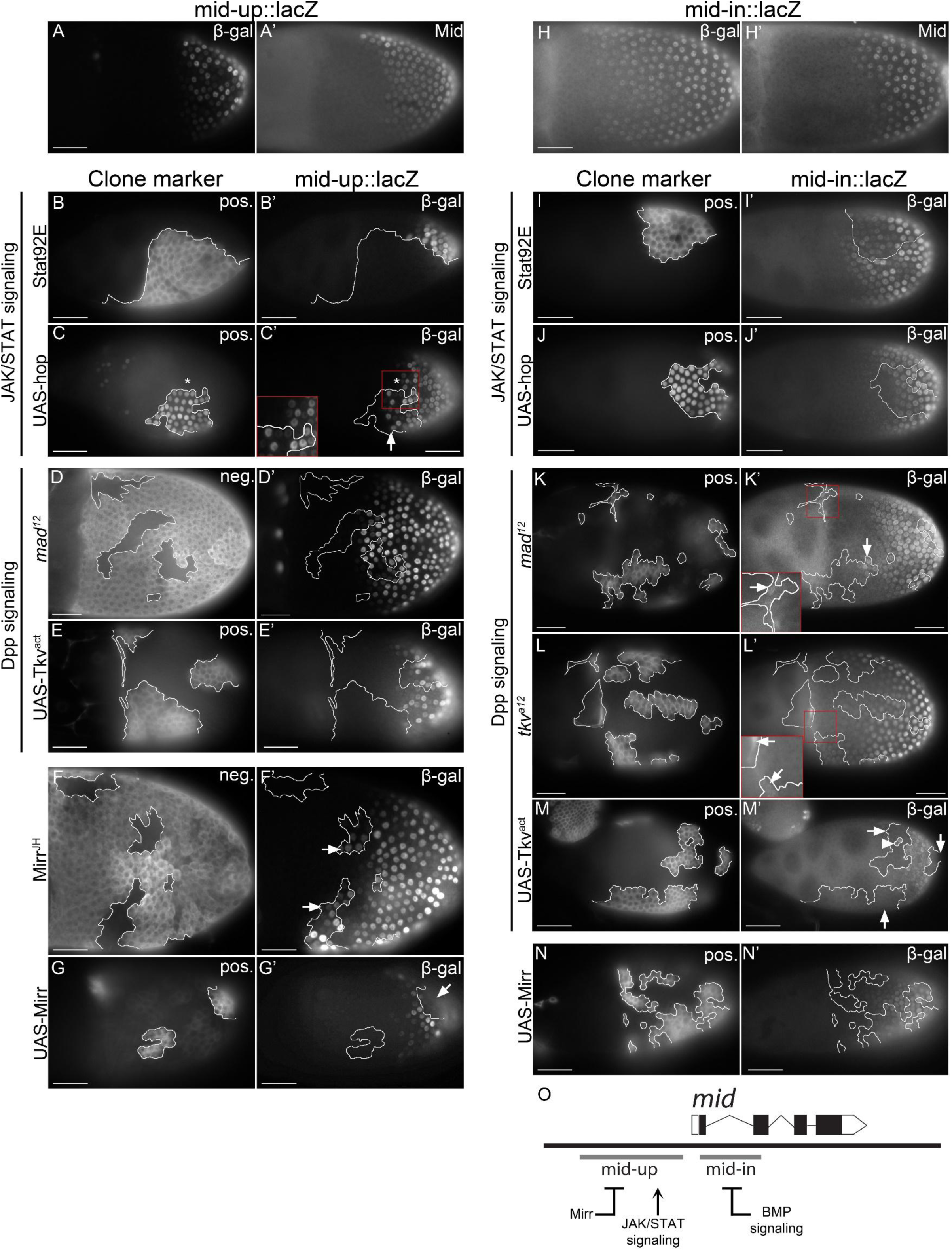
Analysis of the response of mid-up and mid-in reporters to known regulatory inputs. Expression of mid-up and mid-in lacZ reporters when JAK/STAT signaling, Dpp signaling, or Mirr level is altered in clones. White lines mark boundaries of each clone. Scale bars = 50μm. (A-A’) Wild type mid-up lacZ reporter expression for comparison. (B-C’) mid-up reporter expression (β-gal) in JAK/STAT signaling loss-of function (*StatS2E*) or gain-of-function (UAS-*hop*) clones, positively marked by GFP. mid-up reporter expression is lost in *StatS2E* clones (B’, 25/25 posterior clones showed reduced or loss of expression) and ectopically expressed in posterior UAS-*hop* clones (C’, arrow, 10/17 posterior clones showed increased expression, 7/17 showed no change; 4/7 dorsal anterior clones showed increased or ectopic expression, 3/7 showed no change). (D-E’) mid-up reporter expression in BMP loss-of function (*Mad^12^*) or gain-of function (UAS-*tkv^act^*) clones, negatively marked by the absence of NM or positively marked by GFP, respectively. mid-up reporter expression is unchanged in *Mad^12^* clones (D’, 26/36 clones showed no change, 5/36 showed induction; 5/36 showed loss) or in UAS-*tkv^act^* clones (E’, 37/49 clones showed no change; 5/49 showed induction, 7/49 showed loss). In both cases, half of the responding clones show increased expression and half show decreased expression, with no obvious correlation with position or stage. This lack of a clear trend suggests that the apparent effect in some clones is coincidental, due to the cell-to-cell variability of mid-up reporter expression. (F-G’) mid-up reporter expression in *mirr* loss-of function (*mirr^JH^*) and gain-of function (UAS-*mirr*) clones, negatively marked by the absence of NM and positively marked by GFP respectively. mid-up reporter expression is expanded anteriorly in *mirr^JH^* clones (F’, arrows, 34/57 clones at the expression margin showed anterior expansion. 1/57 showed reduction, 22/57 showed no change in expression) and lost in UAS-*mirr* clones (G’, arrow, 24/39 clones showed loss of expression; 0/39 showed induction, 15/39 showed no change). (H-H’) Wild type mid-in lacZ reporter expression for comparison. (I-J’) mid-in reporter expression in *StatS2E* or UAS-*hop* clones positively marked by GFP. mid-in reporter expression is unchanged in *StatS2E* clones (I’, 18/20 clones within reporter expression domain showed no change, 2/20 showed reduced expression) or UAS-*hop* clones (J’, 16/19 clones within reporter expression domain showed no change, 2/19 showed induction, 1/19 showed reduced expression). (K-M’) mid-in reporter expression in BMP loss-of function (*Mad^12^*, *tkv^a12^*[40]) or gain-of function (UAS-*tkv^act^*), positively marked by GFP. (K) mid-in reporter expression is expanded towards the anterior in most *Mad^12^* mutant clones (K’ arrow, 11/16 clones at the dorsal anterior border of reporter expression domain showed induction, 5/16 showed no change; 3/5 Centripetal cell clones showed induction) and *tkv^a12^* mutant clones (L’ arrow, 12/16 clones at the dorsal anterior border of reporter expression domain showed induction, 4/16 showed no change2/3 centripetal cell clones showed induction), and lost in UAS-*tkv^act^* clones (M’ arrows, 32/36 clones showed loss or reduction, 4/36 showed no change). (N-N’) mid-in reporter expression in *mirr* gain-of function (UAS-*mirr*) clones, positively marked by GFP. mid-in reporter expression is unchanged in UAS-*mirr* clones (N’, 16/20 clones showed no change; 2/20 showed induction, 2/20 showed loss). (O) Summary of the distribution of regulatory inputs to the putative *mid* CRMs

Conversely, ectopic JAK/STAT signaling [22] leads to ectopic mid-up reporter expression (Figure 4C-C’). Importantly, like endogenous Mid (Figure S4B’), this ectopic mid-up reporter expression is only observed in clones in the posterior and dorsal anterior domains, where Grk/EGFR signaling is also present; this localized response is consistent with our previous observation that Upd itself is not sufficient to induce Mid expression but rather regulates the outcome of EGFR signalling [4]. These results suggest that the genomic region present in mid-up captures sequence elements that respond to Upd in a pattern that resembles that of endogenous Mid. Since all cells contain the reporter but only certain cells respond, this analysis also reveals in which cells the regulatory element(s) contained in the mid-up genomic sequence can be activated by Upd signaling.

We also asked whether mid-up captures the repression of endogenous *mid* by Dpp signaling, speculating that such negative regulation contributes to the restriction of mid-up expression to the posterior domain. Instead, expression of mid-up reporter is unaffected in homozygous clones for an amorphic allele of the Dpp pathway effector *Mad* [24] (Figure 4D-D’), unlike endogenous *mid* expression, which expands anteriorly in dorsal anterior follicle cell clones (Figure S4C’) [4]. Conversely, ectopic Dpp signaling [23], results in loss of endogenous Mid expression, but has no effect on mid-up reporter expression (Figure 4E-E’, Figure S4D’). These results show that mid-up does not contain sequence elements that respond to Dpp signaling.

Endogenous Mid expression is also repressed by Mirr, refining the anterior limit of the Mid domain in early stages [3], and this negative regulation is also seen with mid-up. Like endogenous Mid, the mid-up reporter is ectopically expressed in cells homozygous for a null allele of *mirr* (Figure 4F-F’, Figure S4E’) and lost in cells expressing ectopic Mirr (Figure 4G-G’, Figure S4F’). These data show that mid-up reporter expression is repressed by Mirr.

### mid-in responds to Dpp, but not to Upd or Mirr

Using the same approach, we found that, unlike mid-up, mid-in reporter expression is regulated by Dpp but not Upd or Mirr. Dorsal anterior clones homozygous for null alleles of *Mad* or *tkv,* and adjacent to the mid-in reporter expression domain, exhibit mid-in reporter expression beyond its normal anterior limit, visible especially between stage 9 and 10B (Figure 4K-L’); this spatiotemporal bias is consistent with the requirement for Grk/EGFR signaling in *mid* expression. Loss of *Mad* and *tkv* function also leads to ectopic mid-in reporter expression in centripetal cells (Figure 4K’, L’). Conversely, clonal activation of Dpp signaling leads to loss of mid-in reporter expression in clones throughout the mid-in expression domain, except those in the posterior-most region (Figure 4M-M’), suggesting that mid-in contains a *cis*-regulatory element that mediates repression by Dpp.

Unlike mid-up or endogenous Mid, mid-in does not respond to Upd or Mirr. Expression of mid-in reporter is unaffected in clones lacking *StatS2E* function (Figure 4I-I’). Consistent with this observation, ectopic JAK/STAT signalling does not induce mid-in reporter expression (Figure 4J-J’). Similarly, mid-in does not respond to ectopic Mirr (Figure 4N-N’). The response of endogenous Mid to these manipulations serves as internal controls (Figure S4).

### Concatenation of mid-up and mid-in does not reconstitute endogenous *mid* expression pattern and result in an additive pattern

Since the responses to Dpp signaling and to Mirr are distributed across the mid-up and mid-in CRMs, we asked whether a single reporter bearing both the mid-upand mid-in sequences would reconstitute the endogenous *mid* regulation and expression pattern. We created two versions of a “fused” *mid* CRM where mid-upand mid-in are placed together upstream of lacZ: one directly fusing the two sequences (mid-[up+in]) and the other containing an intervening 767bp lambda phage-derived sequence that recreates the endogenous spacing between the mid-up and mid-in regions (mid-[up+λ+in]) (Figure 5A). Rather than reconstituting the endogenous *mid* expression pattern, the pattern of both mid-[up+in] and mid-[up+λ+in] reporter expression appears additive: it progressively expands anteriorly similar to mid-in, and it exhibits variable expression levels at the posterior similar to mid-up (Figure 5B-K). This expression pattern suggests that enhancer activities of these regions act additively when concatenated in this transgenic reporter context.

**Figure 5.**
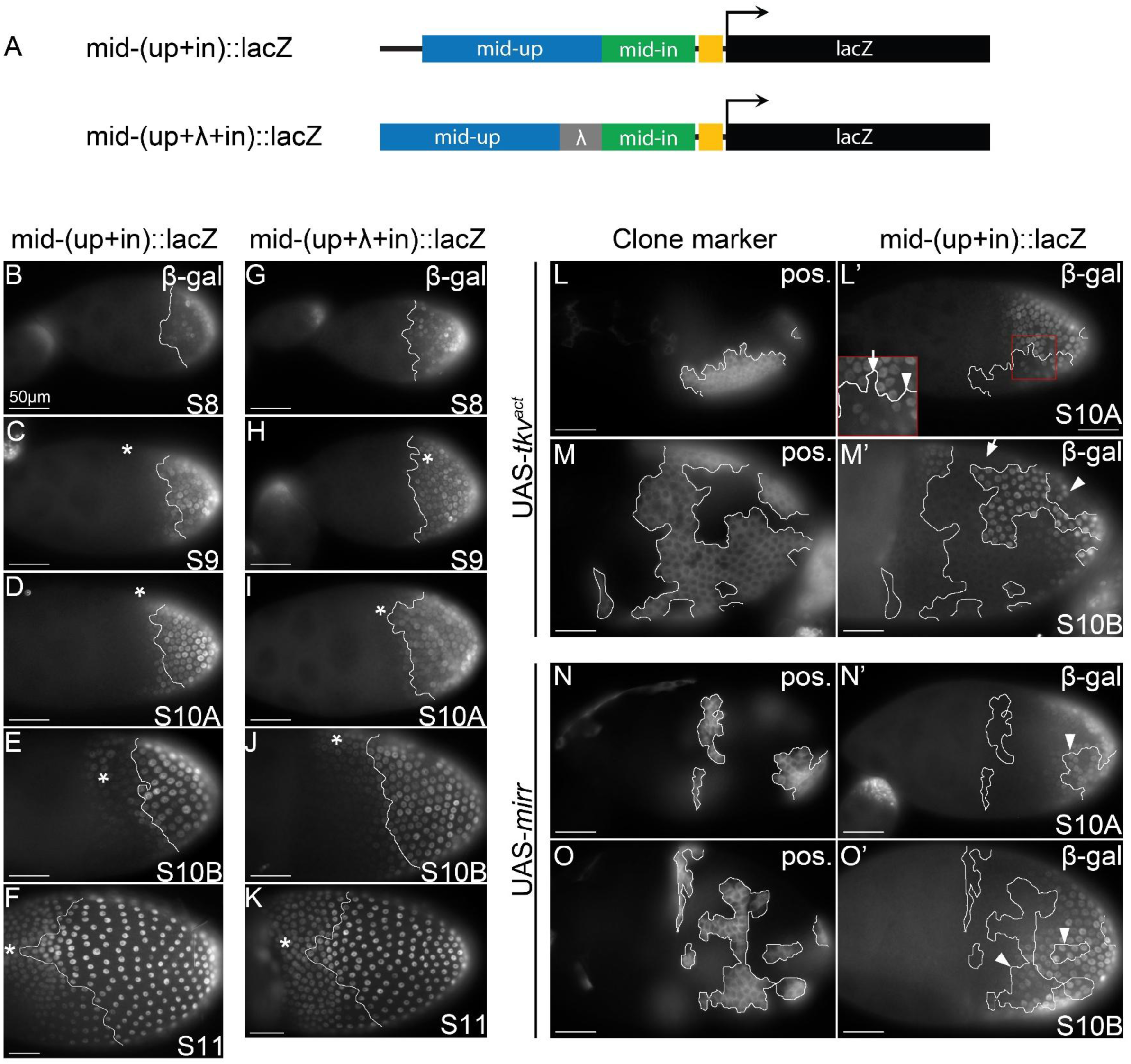
mid-[up+in] reporter expression. (A) Schematic of the mid-[up+in] and mid[up+λ+in] reporters. The mid-up and mid-in genomic regions are concatenated in the mid-[up+in] reporter. mid-up and mid in are separated by a 767bp spacer sequence from lambda phage DNA (λ) in the mid-[up+λ+in] reporter. (B-K) Expression of mid-[up+in] and mid-[up+λ+in] reporters at different developmental stages (S8-S11). White lines mark the expression boundary of the endogenous Mid domain and asterisk marks the dorsal side of the egg chamber where applicable. mid-[up+in] and mid-[up+λ+in] drive reporter expression in posterior follicle cells which expand anteriorly as development progresses, similar to mid-in and unlike mid-up or Mid. (L-M’) mid-[up+in] reporter expression in BMP gain of function clones, positively marked by GFP. White lines mark boundaries of each clone. Expression of mid-[up+in] reporter is lost or reduced in UAS-*tkv^act^* clones within reporter expression domains (arrow) less so in posterior clones (arrowhead)(16/24 posterior clones showed loss; 26/26 clones at anterior of reporter expression domain showed loss). (N-O’) mid-[up+in] reporter response to ectopic Mirr. White lines mark UAS-*mirr* clone borders. Expression of mid-[up+in] reporter is not lost in UAS-*mirr* clones (arrowheads, 28/28 clones within reporter expression domain showed no changes).

Consistent with this interpretation, the response of the mid-[up+in] reporter to Dpp signaling and to Mirr also appears to be a composite of the mid-up and mid-in responses. Unlike mid-in alone, which is repressed by ectopic Dpp signaling in all columnar follicle cells, repression of mid-[up+in] is detectable only in the anterior domain, likely because the mid-in sequences present in the hybrid reporter remain insensitive to repression and continue to drive expression in the posterior region; supporting this interpretation, levels of this posterior expression are variable, resembling the mid-up reporter (Figure 5L-M’).

Similarly, unlike mid-up alone, mid-[up+in] reporter expression is not lost in posterior cells expressing ectopic Mirr, likely because loss of the “mid-up contribution” to reporter expression is obscured by the continuing presence of the mid-in contribution (Figure 5N-O’).

These observations suggest that the mid-up and mid-in CRMs function independently, even when concatenated in the mid-[up+in] reporter, implying that additional sequence context, such as hidden silencers and insulators or chromatin context, is missing when these regions are isolated in the transgenic reporter context.

### mirr-early is regulated by Upd, but not by Dpp or Mid

In early stages of oogenesis, endogenous *mirr* expression is repressed by Upd in the terminal regions of the follicular epithelium, confining *mirr* expression to the intervening “main body” region [6, 7]. The mirr-early reporter exhibits a similar pattern (Figure 6A-A’) and is also negatively regulated by Upd. Cells lacking the Upd downstream transcription factor Stat92E display ectopic mirr-early reporter expression (Figure 6B-B’), while ectopic JAK/STAT signaling represses mirr-early reporter expression (Figure 6C-C’)[4]. These data show that mirr-early contains one or more *cis*-regulatory sequences that respond to repressive input from Upd.

**Figure 6.**
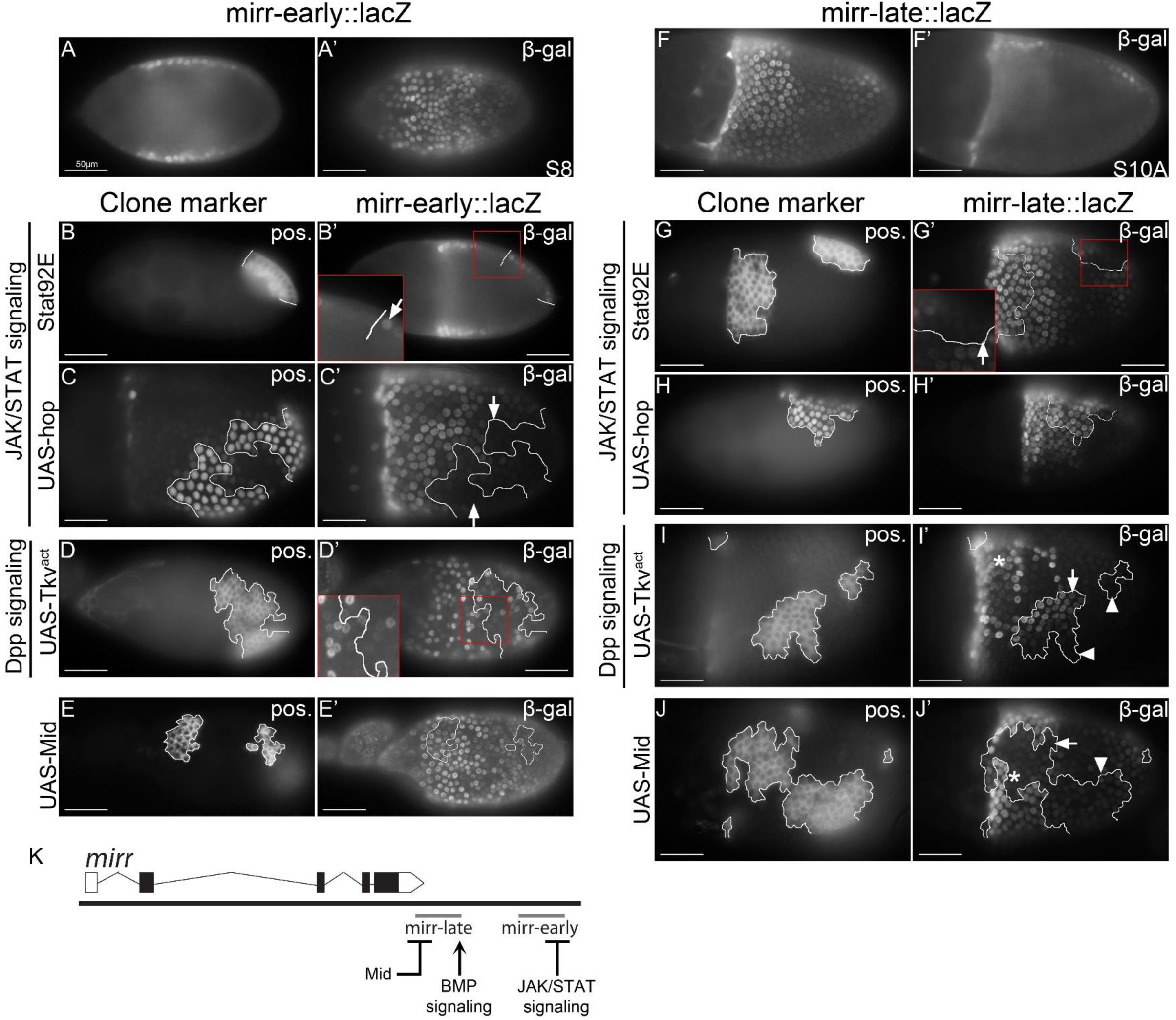
Analysis of the response of mirr-early and mirr-late reporters to known regulatory inputs. Expression of mirr-early and mirr-late lacZ reporters when JAK/STAT signaling, Dpp signaling, or Mid is altered in clones. Clones are positively marked by GFP; white lines mark clone borders. Asterisks mark the dorsal side of the egg chamber where applicable. Scale bars = 50μm. (A-A’) Wild type mirr-early lacZ reporter expression. (B-C’) mirr-early reporter expression (β-gal) in JAK/STAT signaling loss-of function (*StatS2E)* or gain-of-function (UAS-*hop*) clones. mirr-early reporter is ectopically expressed in posterior *StatS2E* mutant clones (B’, arrow, 27/27 clones showed induction) and lost in UAS-*hop* clones (C’, arrows, 25/25 clones showed loss). (D-D’) mirr-early reporter expression in BMP gain-of function (UAS-*tkv^act^*) clones. mirr-early reporter expression is not induced in posterior UAS-*tkv^act^* clones (D’, 43/91 clones showed no change in expression; 14/91 clones showed induction; 34/91 clones showed reduction).E-E’) mirr-early reporter expression in Mid gain-of function (UAS-*mid*) clones. Mirr-early reporter expression is unchanged in UAS-*mid* clones (E’, 19/19 clones showed no change). (F’’) Wild type mirr-late lacZ reporter expression as comparison. (G-H’) mirr-late reporter expression in JAK/STAT signaling loss-of function (*StatS2E*) or gain-of-function (UAS-*hop*) clones. mirr-late reporter expression at the dorsal anterior expression domain is unchanged in *StatS2E* clones, but the expression at the posterior domain is lost in *StatS2E* clones (G’, 19/21 dorsal anterior clones showed no change, 2/21 showed induction, 7/7 dorsal posterior clones showed reduction or loss) or UAS-hop clones (H’, 7/9 dorsal anterior clones showed no change in expression, 2/9 showed induction; 6/8 dorsal posterior clones showed induction, 2/8 showed no change(I-I’) mirr-late reporter expression in BMP gain-of function (UAS-*tkv^act^*) clones. mirr-late reporter expression is induced in UAS-*tkv^act^* clones at the dorsal anterior side (I’, arrow, 24/24 clones at the dorsal anterior showed induction). (J-J’) mirr-late reporter expression in Mid gain-of function (UAS-*mid*) clones. Mirr-late reporter expression is lost in UAS-*mid* clones (J’, arrow, 24/24 dorsal anterior clones showed reduction or loss, 11/11 dorsal posterior clones showed no change; 11/12 ventral clones showed no change, 1/12 showed induction). (K) Summary of the distribution of regulatory inputs to the putative *mirr* CRMs

In contrast to endogenous *mirr*, mirr-early reporter expression is not induced by ectopic Dpp signaling, instead becoming more variable and occasionally even reduced (Figure 6D-D’). This observation suggests that, unlike the endogenous *mirr* locus, mirr-early is not positively regulated by Dpp signaling. This lack of positive regulation is consistent with the observation that the expression pattern of both *mirr* and mirr-early is established earlier than Dpp signaling activity is detected[9].

Finally, mirr-early reporter expression is not repressed by ectopic Mid (Figure 6E-E’, Figure S5). This lack of repression is consistent with the observation that the posterior limit of the mirr-early expression domain overlaps with the anterior limit of the Mid domain (compare Figure 3A-D”’), unlike endogenous *mirr* at these stages, supporting the interpretation that mirr-early does not include regulatory elements that respond to Mid.

### mirr-late responds to inputs that control expression of *mirr*, but the response is complex

The mirr-late reporter is expressed in a complex pattern that includes components of the endogenous *mirr* pattern in later stages as well as a novel posterior expression domain.

Like endogenous *mirr,* which is positively regulated by Grk and Dpp in later stages mirr-late is expressed in the dorsal anterior region of the tissue where these two signals overlap [5, 7, 25, 26]. Consistent with this pattern, ectopic Dpp signaling induces mirr-late reporter expression, but only in the dorsal anterior domain, where Grk is also present, (Figure 6I-I’). These observations suggest that mirr-late contains a BMP responsive element that activates expression in response to Grk/EGFR signaling, analogous to the regulation of endogenous *mirr*.

This anterior expression of the mirr-late reporter also responds to Mid. Clonal expression of ectopic Mid in this region leads to reduced mirr-late expression (Figure 6J-J’, arrow) suggesting that the mirr-late region may contain elements involved in the repression of endogenous *mirr* by Mid. In contrast, this dorsal anterior mirr-late reporter expression is unaffected by either loss of Stat92E function or ectopic JAK/STAT signalling (Figure 6G-H’), suggesting in dorsal anterior cells the mirr-late reporter does not respond to Upd.

As mentioned above, the mirr-late genomic region also drives reporter expression in posterior follicle cells, where endogenous *mirr* is repressed by both Upd and Mid, and therefore not detected. The observation that posterior cells express mirr-late even though they also experience JAK/STAT signaling and express endogenous Mid, suggests that mirr-late is refractory to repression by both of these inputs. Consistent with this prediction, ectopic Mid has no effect on mirr-late expression in these cells (Figure 6J-J’, arrowhead).

Similarly, mirr-late is not repressed but rather positively regulated by Upd. Loss of STAT92E function abolishes posterior mirr-late reporter expression (Figure 6G’), while ectopic JAK/STAT signaling leads to ectopic posterior reporter expression (Figure 6H-H’). These data suggest that mirr-late is expressed in posterior follicle cells because, unlike endogenous *mirr*, it is not repressed by Mid and Upd.

## Discussion

In this study we have identified and characterized two genomic fragments from the *mid* locus (mid-up and mid-in) and two from the *mirr* locus (mirr-early and mirr-late) that drive overlapping patterns of reporter expression in the main-body follicle cells and posterior follicle cells respectively. Multiple enhancers that are associated with the same gene and drive overlapping expression, also known as shadow enhancers, have been identified in a range of developmental contexts [27]. Consistent with their similar expression patterns, such enhancers are often functionally redundant or partially redundant. Indeed, in the case of *mid*, CRISPR-mediated deletion of the endogenous mid-up or mid-in sequence alone has only modest effects on the extent of the Mid expression domain [16]. The presence of such multiple seemingly redundant enhancers is thought to play a role in buffering noise from upstream input, thus conferring robustness of gene expression [27–30].

Despite their similar expression patterns, shadow enhancers from the same locus can be responsive to different regulatory inputs [31]. Our analysis of these *mirr-* and *mid-*associated reporters shows this differential regulation as well, with the two CRMs for each locus responding to a different subset of the known upstream inputs that regulate the endogenous *mirr* and *mid* genes in this tissue [9]. These responses largely align with the reporter expression pattern. For example, mirr-early reporter expression is repressed by JAK/STAT signaling, consistent with its exclusion from anterior and posterior terminal domains , but not by ectopic Mid, consistent with the overlap between the posterior limit of mirr-early expression and the anterior limit of the endogenous Mid domain. Similarly, mirr-late reporter expression in the anterior region of the epithelium is activated by Dpp signaling, while the posterior limit of this domain is defined through repression by Mid. For the *mid* reporters, the observation that mid-up reporter expression is confined to the posterior region is consistent with its positive regulation by JAK/STAT signaling and its repression by Mirr.

An apparent exception to this alignment is the progressive anterior expansion of mid-in reporter expression in later stages, which seems inconsistent with our observation that mid-in is repressed by ectopic Dpp signaling. However, this anterior expansion pattern <u>fits</u> well with the dynamic expression pattern of the Tkv receptor, which is initially present in all follicle cells but gradually recedes toward the anterior as development proceeds [32–34], leaving cells unable to respond to Dpp. This anterior shift in the endogenous Tkv expression pattern would lead to loss of Tkv-mediated repression of mid-in and is thus consistent with the anteriorly expanding mid-in reporter expression.

Another interesting example is the mirr-late reporter expression at the dorsal posterior domain, which is not observed for endogenous *mirr.* This dorsal posterior expression overlaps the endogenous Mid domain, and therefore is not consistent with the observation that mirr-late is repressed by ectopic Mid. The regulation of this mirr-late reporter expression domain is also unexpected, as it is activated instead of repressed by JAK/STAT signaling in the posterior domain. The reason for this emergent expression domain and regulatory relationship is elusive, but such scenarios may reveal interesting aspects of the sequence or cellular context.

Discrepancies like these highlight the important role for sequence context in enhancer function. Genomic regions isolated in a reporter setting may lack important regulatory elements that contribute to patterned expression. For example, the absence of additional relevant enhancers, silencers or boundary elements that are not included in the reporter constructs, or more general mechanisms like chromatin states, can affect promoter activity in a given cellular or tissue context [35]. Consistent with this notion, the apparently additive expression pattern and regulation of the mid-[up+in] reporter suggest that simply incorporating multiple response elements into the same reporter is not sufficient to mimic the endogenous *mid* pattern, implying that relevant sequence elements are lacking.

Transcriptional reporters have a long history as genetic tools for dissecting *cis* regulatory logic and remain a valuable strategy for isolating and visualizing enhancer activity and for testing regulatory relationships *in vivo.* Such a reductionist framework has been particularly valuable for identifying putative CRMs associated with loci, such as developmental genes as mentioned above, that bear multiple partially redundant enhancers. Since the contribution of a given regulatory element may be masked in the endogenous locus by such redundancy, transcriptional reporters can reveal putative regulatory elements whose individual contributions would not have been otherwise detected [27].

More recently, this focused region-by-region strategy of CRM mapping and identification has been complemented by the development of chromatin level assays such as ChIP and ATAC seq as genome-wide approaches to CRM identification that rely on detection of physical interactions as an indication of enhancer activity. However, accumulating evidence suggests that relying on a single approach is not sufficient. For example, a recent analysis found that chromatin level assays do not agree with data from CRM reporter analysis, often failing to identify CRMs that had been identified through reporter analysis [36]. This observation could suggest that these chromatin-level approaches may fail to identify potential functional elements or that, conversely, transcriptional reporters may identify elements whose potential enhancer activity would normally be constrained in the larger sequence or chromatin environment in the endogenous context [37]. This discrepancy speaks to the need for a diverse set of tools to approach the identification and study of enhancers.

While identifying endogenously relevant enhancers is an important goal of CRM identification, even sequence elements that drive novel expression patterns in the context of a transcriptional reporter can be informative. For example, patterns <u>of</u> reporter expression can reveal similarities between cells that might not have been observed previously in known patterns of gene expression or cell fate. Indeed, the novel dorsal posterior domain of mirr-late reporter expression reveals a commonality between these cells that sets them apart from the rest of the cells in the posterior domain – namely their ability to express the mirr-late reporter. In fact, a fundamental goal of the two genome-wide projects that created libraries of transgenic “tile-line” reporters, each bearing a different defined fragment of non-coding DNA [11, 14], was to identify novel neuronal cell populations defined by their ability to express a given reporter. Regardless of their function in the endogenous context, transcriptional reporters like these can be interesting tools for asking whether candidate CRMs can respond directly to upstream inputs. [3] While there are many possible indirect-regulation scenarios, in some cases the reporter could be regulated directly through binding of the effector transcription factors to genomic sequences present in the reporter. In this respect, characterization of the expression and regulation of transcriptional reporters can set the stage for more direct studies of transcriptional regulation during tissue patterning.

## Materials and Methods

### Drosophila Strains

The following Drosophila stocks were used: GAL4 transcriptional reporters (see Table S1 for details; [14, 15]), FRT82B *StatS2E^85CS^* [17]; UAS-*tkv^actQ1SSD^* (i.e. UAS-*tkv^act^*) [23]; UAS-*mid* [38];

UAS-*hop* [22]; *mirr^JH^* FRT80B [4]; *mirr*-*lacZ* (*mirr^l(3)cD1^*) [5]; UAS*-mirr* [39]; *Mad*^12^ FRT40A [24], and *tkv^a12^* FRT40A [40].

The following stocks were used for generating clones:

UAS-GFP, yw hsFlp; tubGAL80 FRT40A; tubGAL4 (RRID:BDSC_605350)

yw hsFlp; tubGal80 piM FRT40A/ CyO; tubGal4 / TM6B, Hu UAS-GFP, hsFlp; tubGal4 FRT82B tubGAL80/ TM6B

yw hsFlp, tubGAL4, UAS-GFP; tubGAL80 FRT80B / TM6B Tb Hu (RRID:BDSC_42732) yw hsFlp; NM FRT80B (RRID:BDSC_1949)

### Transgenic reporter screen

Drosophila strains containing transgenic GAL4 transcriptional reporters, each containing 1-3 kb of genomic DNA placed upstream of an hsp70 basal promotor driving expression of the GAL4 transcriptional regulator were screened for expression in the ovary. We tested lines assigned to the *mirr*, *mid*, and *H15* loci [14, 15]; see Table S1), by crossing them to strains containing a UAS-GFP transgene and assessing their ability to drive ovarian expression of GFP. Select strains exhibiting patterned GFP expression in the follicular epithelium were chosen for further analysis.

### Cloning and transgenesis of lacZ reporter constructs

To generate lacZ versions of each selected GAL4 reporter, the relevant genomic sequences were amplified from genomic DNA and cloned into a linearized placZ.attB vector [41].

These constructs were then integrated into the Drosophila genome using the PhiC31 integrase and the attP landing site at cytological positions 58A or 86Fb. For each transformant strain, genomic DNA was amplified and genotyped by either restriction digestion or sequencing to validate the identity of the transgene.

### Mosaic Analysis

Negatively marked loss-of-function clones were generated as described previously [42] through FLP/FRT-mediated mitotic recombination between a wild-type or mutant chromosome bearing P{ry[+t7.2] = neoFRT} 40A, 80B, or 82B and a marker chromosome bearing the same FRT and a P{w[+mC] = NM} insertion on the same chromosome arm. The MARCM system [43] was used to generate positively marked clones of cells driving expression of either P{UAS-mCD8::GFP.L} or P{UAS-GFP.nls}. In some experiments, the GAL4 expressed in these clones also drove the expression of a UAS-*mirr*, UAS-*mid*, UAS-*tkv^act^*, or UAS-hop transgene. For generation of clones, females of the appropriate genotype were heat shocked for 1 hr at 37°C as newly eclosed adults, aged at 22°C for 7-10 days, then supplemented overnight with dry yeast before dissection to harvest ovaries.

### Immunohistochemistry

Immunostaining was carried out as described previously [42]). Antibodies used were guinea pig anti-Mid (1:2,000) [3], rabbit anti-β-gal (1:10,000, MP Biomedicals-Cappel), mouse anti-β-gal hybridoma supernatant 40A-1A-s (1:200, Developmental Studies Hybridoma Bank), rabbit anti-c-Myc supernatant sc-789 (1:200, Santa Cruz Biotechnology), and rabbit anti-GFP A6455 (Life Technologies). All secondary antibodies were highly cross-adsorbed Alexa Fluor-conjugated anti-IgG (1:2,000, Invitrogen). All secondary antibodies, as well as anti-Mid, were diluted 1:10 in PBS and pre-adsorbed by incubation with fixed wild-type ovaries or embryos prior to use.

### Microscopy and Image Analysis

Images were acquired using a Zeiss Axioplan 2 wide-field fluorescence microscope. Apart from minor uniform linear adjustments to brightness and contrast, no image manipulations were performed.

## Acknowledgements

We thank Erika Bach, William Brook, Doug Harrison, and Laurel Raftery for fly strains, and Daniella Ansong, Kaitlin Mac Donald, and Gary Zhou for technical assistance. We also thank Kaitlin Mac Donald for helpful comments on the manuscript. Stocks obtained from the Bloomington Drosophila Stock Center (NIH P40OD018537) were also used in this study. This work was funded by the Natural Sciences and Engineering Research Council of Canada.

## Notes

### Competing Interest Statement

The authors have declared no competing interest.

### Summary of Updates

Abstract and Discussion revised; Fixed Typos; Fixed missing scale bars in Figures 4 and 6

