## Supplemental Figures for "Identification of *cis*-regulatory modules responding to positional cues regulating the patterned expression of the *Drosophila* follicle cell fate determinants *mid* and *mirr*"

| Transgene | Locus | Gal4 expression in ovary |
|---|---|---|
| 29787 | mirr | None |
| 29789 | mirr | Germarium |
| 29790 | mirr | FCs, stage 13, dorsal appendages |
| 29791 | mirr | Germarium |
| 29792 | mirr | Germarium; FCs, stage 13, dorsal appendages |
| 29794 | mirr | None |
| 29795 | mirr | None |
| 29796 | mirr | None |
| 29797 | mirr | None |
| 29798 | mirr | None |
| 29799 | mirr | None |
| 29800 | mirr | Germarium |
| 29801 | mirr | None |
| 29804 | mirr | None |
| 29807 | mirr | None |
| 29810 | mirr | None |
| 29811 | mirr | FCs, stages 8-14, enriched in dorsal anterior |
| 29812 | mirr | FCs, stage 13, dorsal appendages |
| 29813 | mirr | None |
| 29814 | mirr | FCs, stages 7-14, restricted from termini |
| 29815 | mirr | FCs, stage 13, dorsal appendages |
| 29816 | mirr | None |
| 29817 | mirr | None |
| 29818 | mirr | None |
| 29820 | mirr | Germarium |
| 29821 | mirr | None |
| 29822 | mirr | None |
| 29823 | mirr | None |
| R34C05 | mirr | None |
| R34D08 | mirr | None |
| R34C01 | mirr | None |
| R34E06 | mirr | None |
| R33B08 | mirr | None |
| R34D12 | mirr | None |
| R34B11 | mirr | FCs, stage 13, dorsal appendages |
| R33B03 | mirr | None |
| R32G07 | mirr | None |
| R33C10 | mirr | FCs, stages 7-14, irregular |
| R33E04 | mirr | None |

| Transgene | Locus | Gal4 expression in ovary |
|---|---|---|
| R34C07 | mirr | None |
| R85B05 | H15 | None |
| R85A11 | H15 | None |
| R85A09 | H15 | None |
| R85A08 | H15 | None |
| R85A03 | H15 | None |
| R85B01 | H15 | None |
| R85A12 | H15 | None |
| R85A10 | H15 | FCs, stage 9, border cells |
| R85B03 | H15 | None |
| R85B04 | H15 | None |
| R85A07 | H15 | None |
| R81H10 | H15 | None |
| R85B06 | H15 | FCs, stage 13, dorsal appendages |
| R85B02 | H15 | FCs, stage 9, border cells |
| R85A02 | H15 | None |
| R85A05 | H15 | None |
| R81H11 | H15 | None |
| R85A01 | H15 | None |
| R85A04 | H15 | None |
| R86G06 | mid | None |
| R86G04 | mid | FCs, stage 7-14, posterior |
| R86F11 | mid | FCs, stage 7-14, posterior |
| R86G05 | mid | None |
| R86F08 | mid | None |

### Table S1. Expression of transgenic GAL4 reporters assigned to the *mirr*, *H15* and *mid* loci

[1] Drosophila strains containing GAL4 *cis*-regulatory reporters analyzed for ovarian expression [14, 15].

[2] Predicted locus assigned to the genomic region contained in the reporter

[3] Detectable GAL4-mediated expression of UAS-GFP in the follicle cells (FC), with details where appropriate[14, 15].

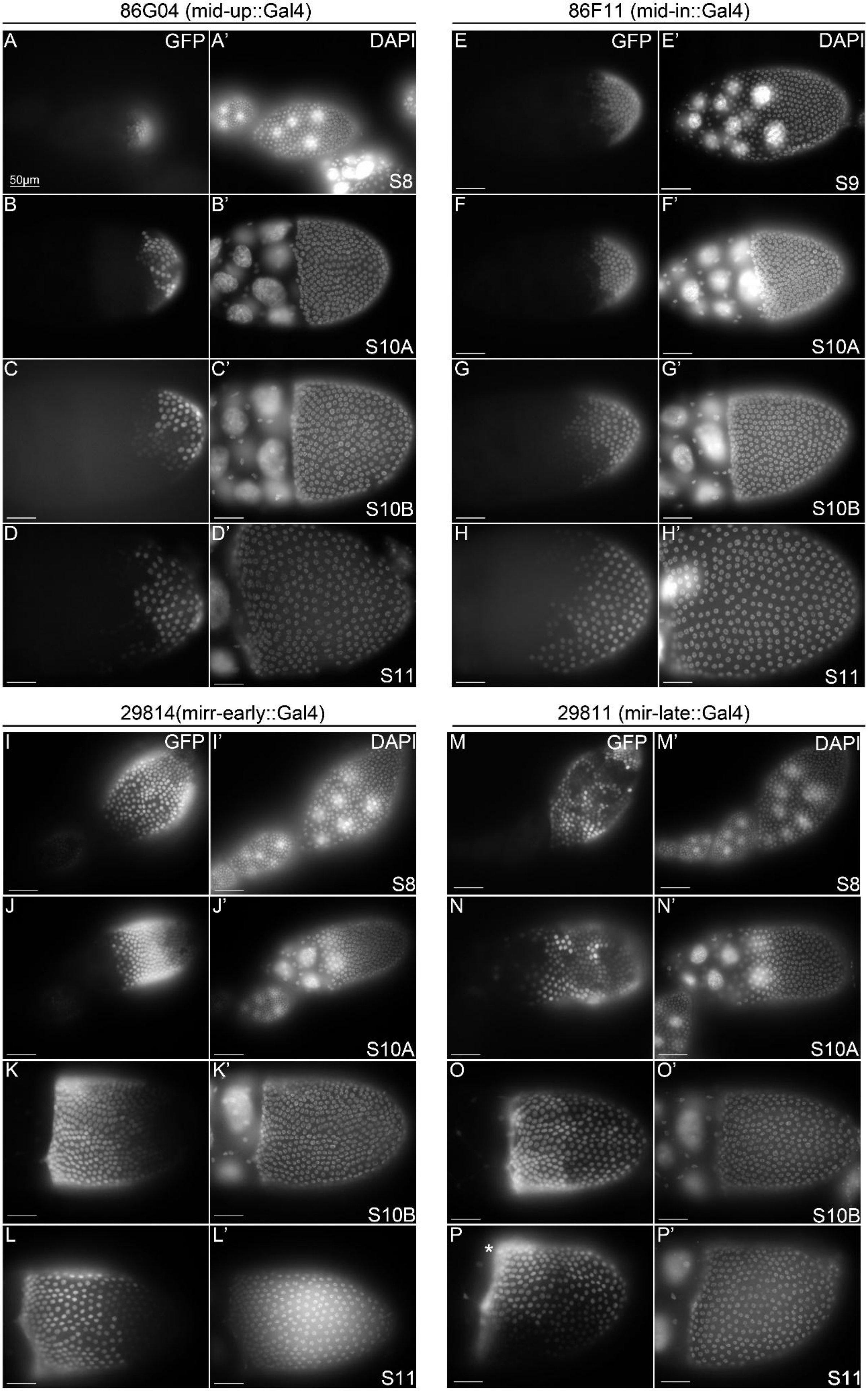

86G04 (mid-up::Gal4)
86F11 (mid-in::Gal4)
A
GFP
A'
DAPI
50μm
S8
E
GFP
E'
DAPI
S9
B
B'
S10A
F
F'
S10A
C
C'
S10B
G
G'
S10B
D
D'
S11
H
H'
S11
29814(mirr-early::Gal4)
29811 (mir-late::Gal4)
I
GFP
I'
DAPI
S8
M
GFP
M'
DAPI
S8
J
J'
S10A
N
N'
S10A
K
K'
S10B
O
O'
S10B
L
L'
S11
P
*
P'
S11

### Figure S1 *mirr* and *mid cis*-regulatory GAL4 reporters closely resemble the lacZ reporters expression patterns.

Putative *mid* and *mirr* GAL4 reporter expression in the follicular epithelium at different developmental stages (S8-11). For each. the GAL4 version of the reporter drives UAS-GFP expression in a similar pattern as the lac-Z version, except that the GAL4 version of mid-in exhibits a more modest late-stage anterior expansion than the lacZ version. Asterisks indicate dorsal position where applicable. Scale bars = 50µm.

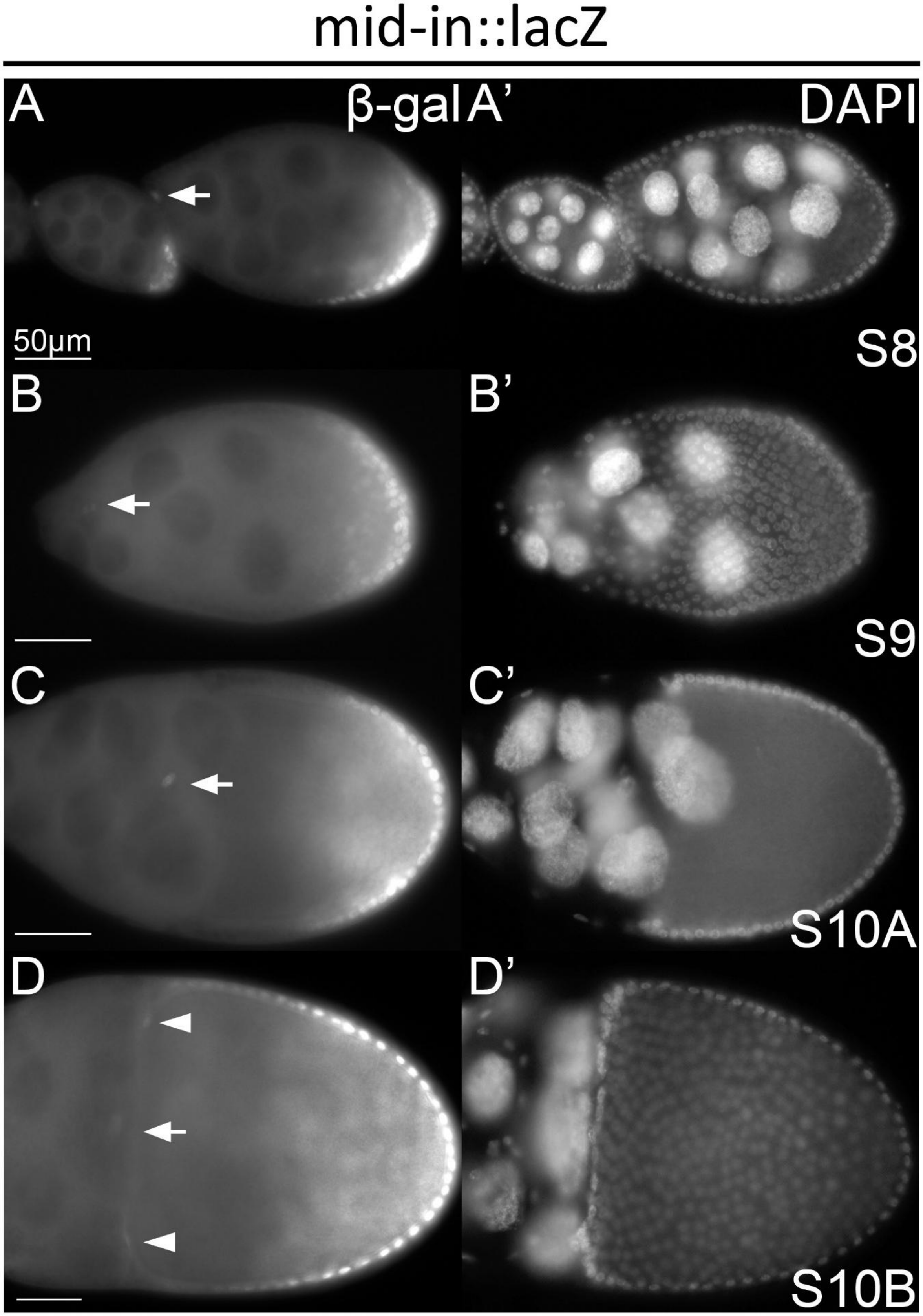

mid-in::lacZ
A
β-gal
A’
DAPI
50μm
S8
B
B’
S9
C
C’
S10A
D
D’
S10B

## Figure S2    Expression of mid-in reporter in polar cell and centripetal cells

Additional details related to Fig. 2. Cross sectional view of mid-in lacZ reporter expression in developing egg chambers. Weak expression of mid-in reporter is detected in the border cell cluster (white arrows) and centripetal cells (arrowhead).

*mirr*$^{l(3)6D1}$ lacZ

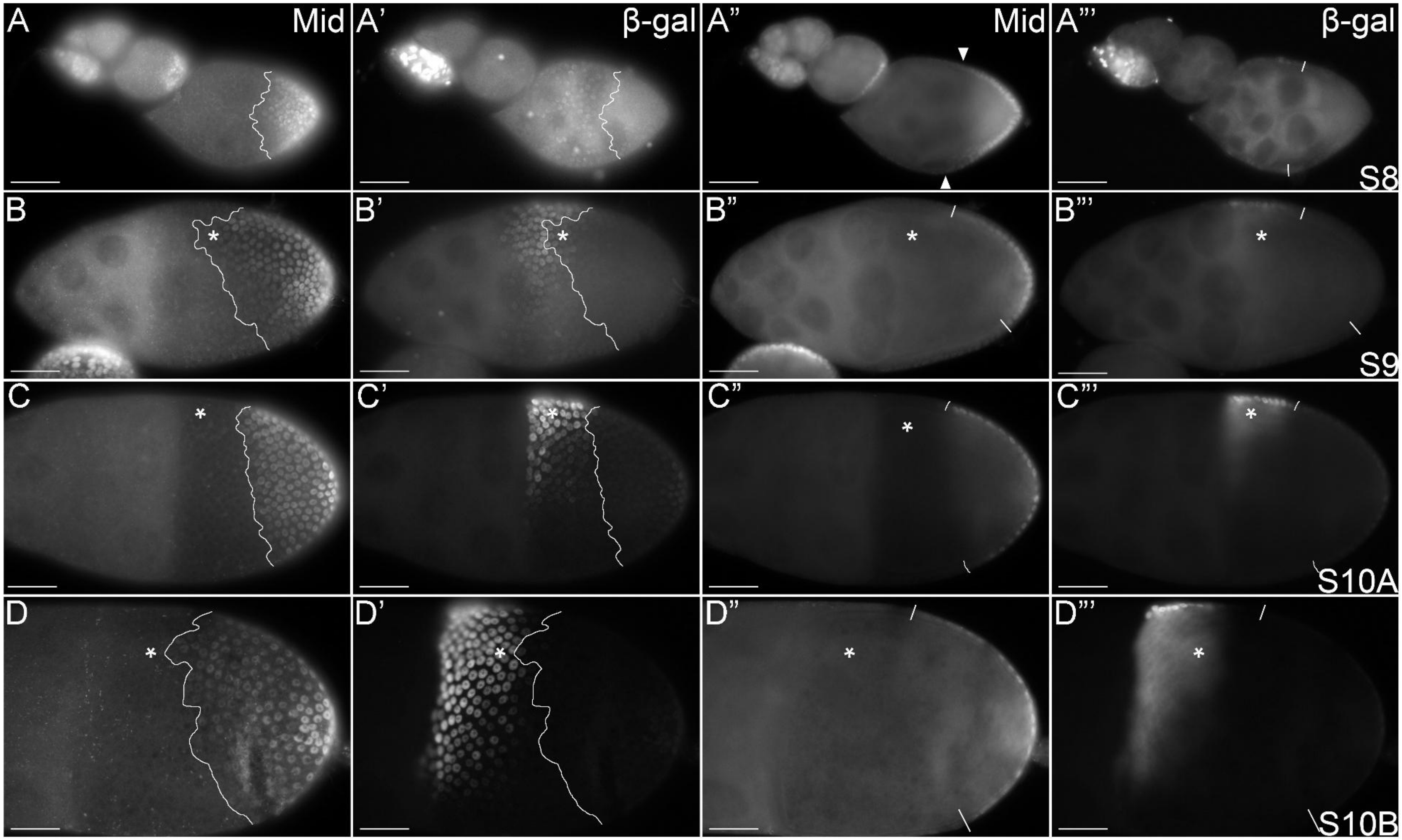

### Figure S3 Expression of the mirr-late regulatory reporter resembles the pattern of endogenous mirr in late oogenesis.

Endogenous expression of *mirr* in stage8-10B egg chambers as observed with an enhancer trap (*mirrl(3)6D1*). The left two columns show the surface view of *mirrl(3)6D1* expression at the follicular epithelium with Mid for comparison; the right two columns show the cross-sectional view of the same egg chambers. White lines mark the expression domain of endogenous Mid, and asterisk marks the dorsal side of the egg chamber where applicable. Scale bar = 50μm. Dorsal anterior regions are complementary and separate from the posterior expression domain of Mid.

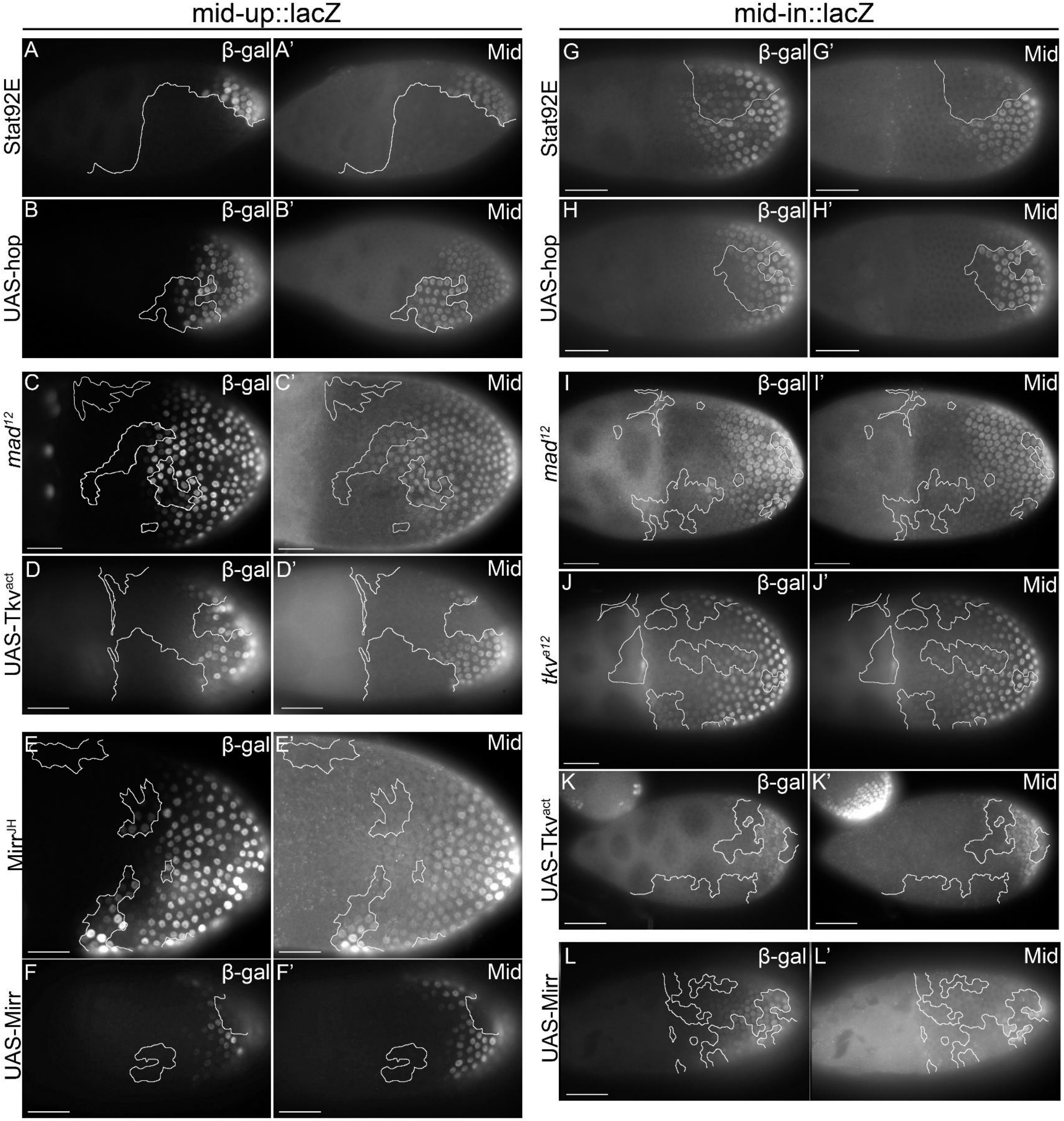

mid-up::lacZ
mid-in::lacZ
A
β-gal
A'
Mid
G
β-gal
G'
Mid
Stat92E
B
β-gal
B'
Mid
H
β-gal
H'
Mid
UAS-hop
C
β-gal
C'
Mid
I
β-gal
I'
Mid
mad12
D
β-gal
D'
Mid
UAS-Tkvact
J
β-gal
J'
Mid
tkva12
K
β-gal
K'
Mid
UAS-Tkvact
E
β-gal
E'
Mid
MirrJH
F
β-gal
F'
Mid
UAS-Mirr
L
β-gal
L'
Mid
UAS-Mirr

### Figure S4 Effect of clonal manipulations of signaling inputs on mid-up and mid-in reporter expression, with levels of endogenous Mid as an internal control

**Internal controls for Fig. 4**. Expression of mid-up and mid-in lacZ reporters when JAK/STAT signaling, Dpp signaling, or Mirr level is altered in MARCM clones. White lines mark boundaries of each mosaic clones.

(A-B') mid-up reporter (A, B) and endogenous Mid (A', B') expressions in JAK/STAT signaling loss-of function (Stat92E) or gain-of-function (UAS-hop) mosaic clones.

(C-D') mid-up reporter (C, D) and endogenous Mid (C' D') expressions in BMP loss-of function (*Mad*$^{12}$) or gain-of function (UAS-*tkv*$^{act}$) mosaic clones.

(E-F') mid-up reporter (E, F) and endogenous Mid (E', F') expressions in Mirr loss of function (*mirr*$^{JH}$) and gain-of function (UAS-*mirr*) mosaic clones

(G-H') mid-in reporter (G, H) and endogenous Mid (G', H') expressions in JAK'/STAT signaling loss-of function (Stat92E) or gain-of-function (UAS-hop) mosaic clones.

(I-K') mid-in reporter (I, J, K) and endogenous Mid (I', J', K') expressions in BMP loss-of function (*Mad*$^{12}$) or gain-of function (UAS-*tkv*$^{act}$) mosaic clones.

(L-L') mid-in reporter (L) and endogenous Mid (L') expressions in Mirr gain-of function (UAS-Mirr) mosaic clones

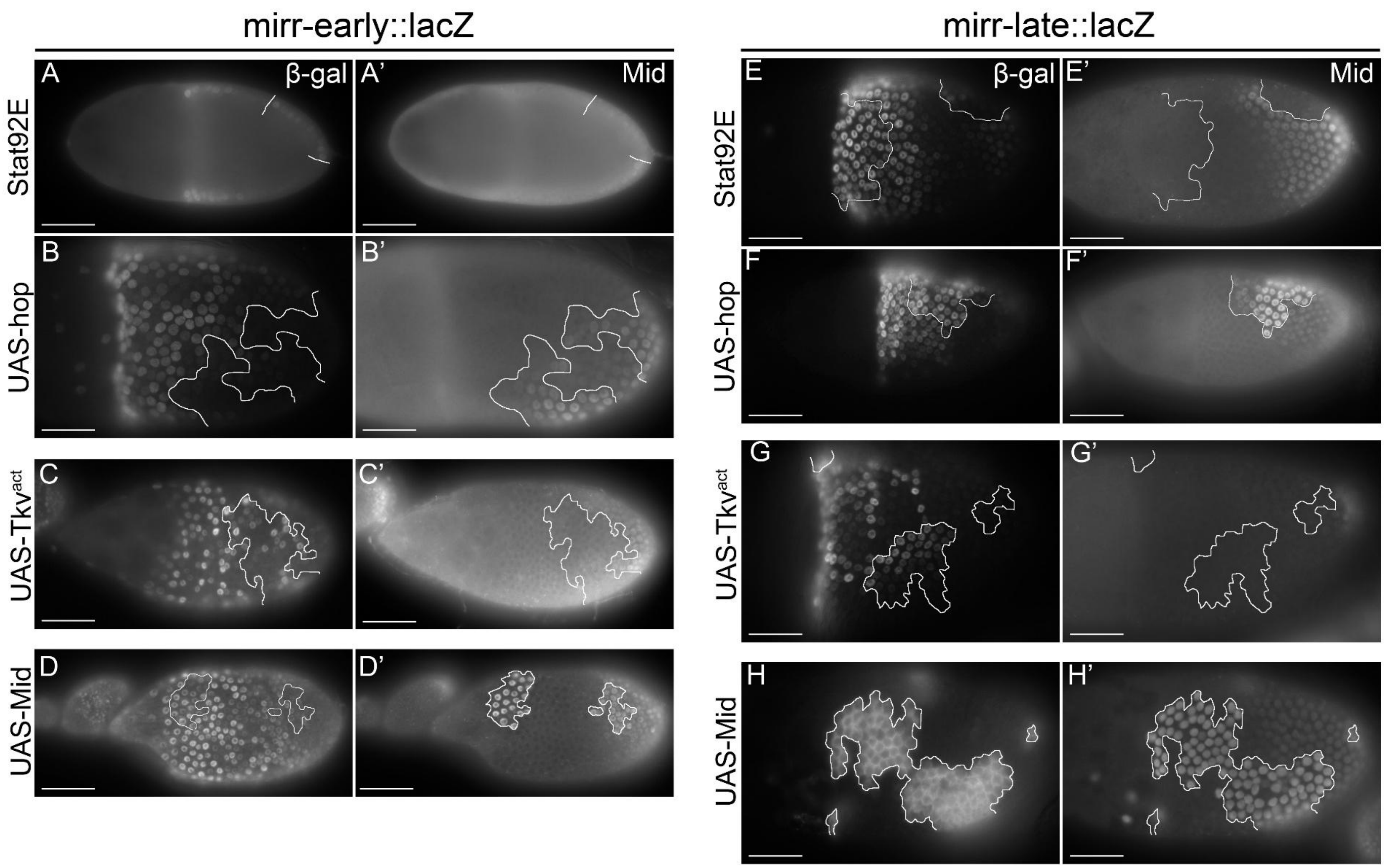

mirr-early::lacZ
mirr-late::lacZ
Stat92E
UAS-hop
UAS-Tkv$^{act}$
UAS-Mid
A
β-gal
A'
Mid
B
B'
C
C'
D
D'
E
β-gal
E'
Mid
F
F'
G
G'
H
H'

### Figure S5 Effect of clonal manipulations of signaling inputs on mirr-early and mirr-late reporter expression, with levels of endogenous Mid as an internal control

**Internal controls for Fig. 5.** Expression of mirr-early and mirr-late lacZ reporters when JAK/STAT signaling, Dpp signaling, or Mid level is altered in MARCM clones. White lines mark clone borders.

(A-B') mirr-early reporter (A, B) and endogenous Mid (A', B') expressions in JAK/STAT signaling loss-of function (Stat92E, UAS-Dome$^{RNAi}$) or gain-of-function (UAS-hop) clones.

(C-C') mirr-early reporter (C) and endogenous Mid (C') expressions in BMP gain-of function (UAS-*tkv*$^{act}$) clones.

(D-D') mirr-early reporter (D) and endogenous Mid (D') expressions in Mid gain-of function (UAS-Mid) clones

(E-F') mirr-late reporter (E, F) and endogenous Mid (E', F') expressions in JAK/STAT signaling loss-of function (Stat92E) or gain-of-function (UAS-hop) clones.

(G-G') mirr-late reporter (G) and endogenous Mid (G') expressions in BMP gain-of function (UAS-*tkv*$^{act}$) clones.

(H-H') mirr-late reporter (H) and endogenous Mid (H') expressions in *mid* gain of function (UAS-Mid) clones.
